# Copper nanoparticles prompt the activity of estuarine denitrifying bacterial communities at relevant environmental concentrations

**DOI:** 10.1101/2020.07.05.188334

**Authors:** Joana Costa, António G.G. Sousa, Ana Carolina Carneiro, Ana Paula Mucha, C. Marisa R. Almeida, Catarina Magalhães, Mafalda S. Baptista

## Abstract

Effects of metallic nanoparticles (NPs) to the estuarine biota have mostly been shown for concentrations higher than those actually measured or predicted in these environments. To address this gap, a range of concentrations expected to occur in estuarine environments (from 0.01 to 1 μg g-1) was employed in microcosms studies to assess the impact of Cu NPs in the denitrification pathway. That was achieved by quantifying gene expression and the potential denitrification rate in estuarine sediments exposed to Cu NPs for up to six days. Expression of nitrite (*nirS*) and nitrous oxide (*nosZ*) reductase genes was enhanced in a timewise manner. For the highest Cu NPs (1 μg g^-1^) an increase in gene expression could be seen immediately after 1 h of exposure, and continuing to be enhanced up until 7 h of exposure. For the lowest Cu NPs (0.01 μg g^-1^) an increase in gene expression could only be seen after 4 h or 7 h of exposure; however it continued to rise up until 24 h of exposure. In any case, after 48 h the expression levels were no longer different from the non-exposed control. Concomitantly to increased gene expression the potential denitrification rate was increased by 30 %. Our results suggest that deposition and adsorption of Cu NPs to estuarine sediments promotes the immediate and transient expression of key genes of the denitrification pathway. The long term impact of continuous inputs of Cu NPs into estuaries deserves renewed analysis to account for their effects, not just on the biota, but especially on ecosystems services.

**Environmental significance:** Interactions of metallic nanoparticles with microbial communities of estuarine sediments are poorly characterized and its impact towards ecosystem services even less. By assessing the effect of copper nanoparticles on the expression of key genes of the denitrification pathway, an essential step for nitrogen (N) removal, we were able to show that denitrifying communities are immediately activated after exposure, increasing the denitrification rates in estuaries. The importance of denitrification lies in its release of dinitrogen (N_2_) to the atmosphere but also in the emissions of N_2_O (a potent greenhouse gas). The results obtained in this study gather data that contribute information on the denitrification dynamics in estuaries, invaluable for a timely response to the expected upcoming changes in coastal areas.

**Table of contents:** 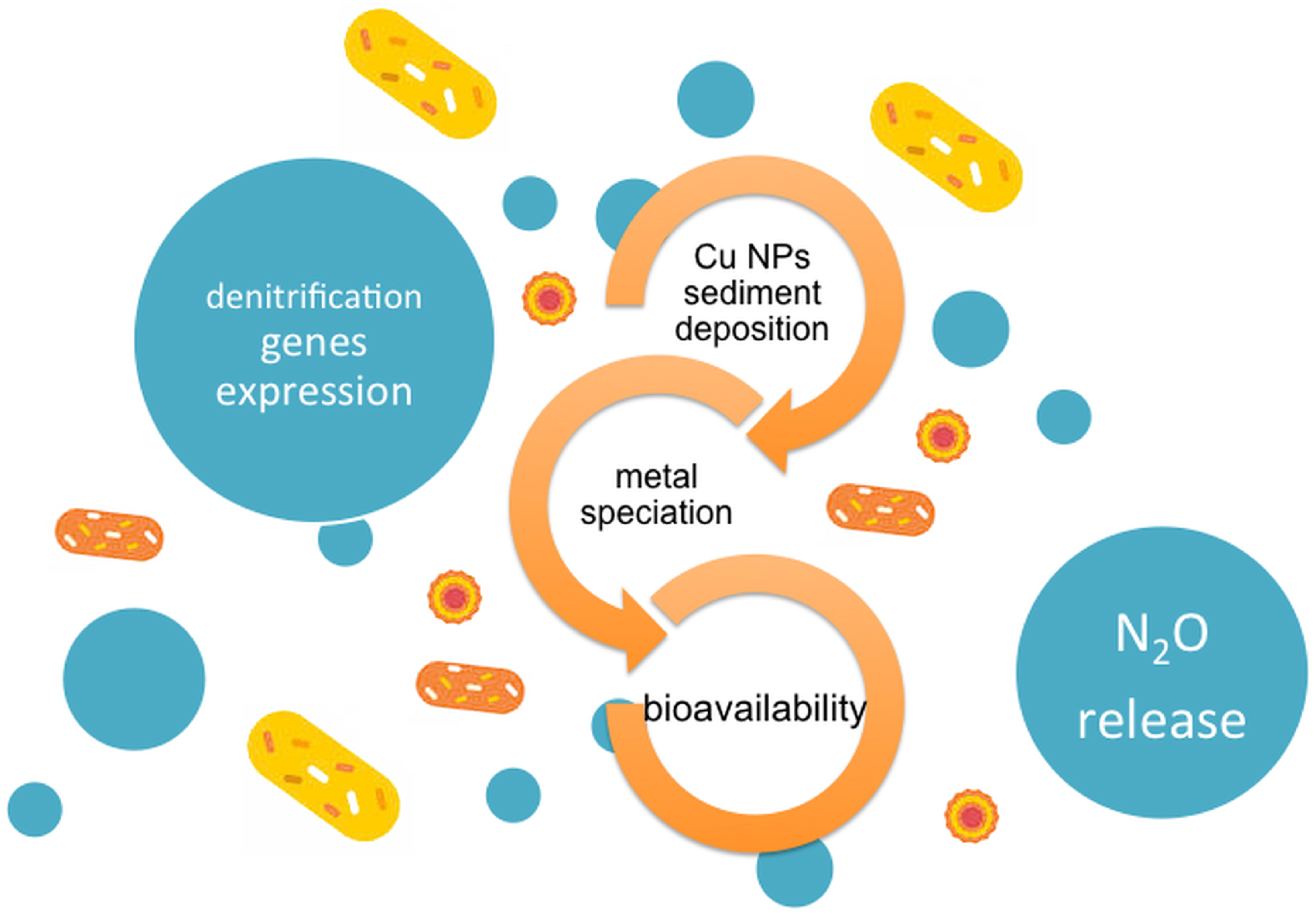

In estuaries the deposition upon the sediments of copper nanoparticles can contribute to change metal availability and promote the activity of denitrifying bacteria

## 1 Introduction

The industrial use of nanoparticles (NPs) has had an exponential growth in the past decades. As a consequence, ultimately the aquatic environment becomes the final destination and repository of these NPs and its impact on the health of the ecosystems is a growing concern in environmental science and management.^1^ One of the major classes of engineered nanomaterials are metallic NPs, such as copper (Cu NPs). Due to their versatility, their applications are broad and Cu NPs are currently used for a variety of purposes such as antimicrobials, treatment drugs, food additives, catalysts, surfactants, sensors and semiconductors.^2^ The vast majority of the environmental Cu NPs are nanoparticles derived from agriculture^3^ and antifouling paints.^4^ In agriculture Cu-based nanomaterials are used as pesticides,^5^ or as biosolids from wastewater treatment plants.^6^ Marine antifouling paints can contain up to 50 % Cu and, therefore, large amounts of Cu can be released and accumulated in coastal areas.^4^

Once in the aquatic environment NPs can undergo transformations such as aggregation, sedimentation, dissolution, sulfidation, amongst others, and can interact with organic matter.^7^ The type of transformation will depend on the size of the NPs, composition of the NPs surface (charge and functionality) and the environmental conditions. In aquatic environments Cu NPs stability is strongly influenced by salinity and pH.^8^ Measurements of Cu NPs dissolution rates have shown this process to be highly variable for freshwater and marine systems.^9^ This variable behaviour poses enhanced difficulties to the assessment of Cu NPs effects, with pronounced differences observed in the responses between species and their physiological variables.^10^ Studies focusing on exposure of aquatic organisms to metallic NPs have shown its consequences to single organisms^11^ but also that, at the community level, species interactions and feedbacks may dampen effects that are seen at the organisms level.^12^ Information related to the environmental risk posed by metallic NPs is still missing largely due to the difficulties involved in the quantification of these particles in the environment.^13^ Because of that, there is great concern in health and environmental issues that may arise from exposure to metallic NPs, even if so far no deleterious effects have been reported, for the range of concentrations expected or predicted to occur in the environment.^14^

In estuaries Cu can greatly impact ecosystem processes. For example, Cu plays a crucial role in denitrification processes since it is a co-factor to NirK NO_2_-reductase and NosZ N_2_O-reductase enzymes.^15^ Previous studies have showed the impact of Cu on the abundance and transcription of genes of the denitrification pathway in an estuarine environment.^16,17^ However, studies specifically addressing the role of Cu NPs (as opposed to ionic Cu), in a range of expected concentrations and of expected sizes are still missing. In this work we address this question by testing the hypothesis that deposition of Cu NPs in estuarine sediments at environmental relevant concentrations and sizes will affect metal bioavailability, which in turn will affect denitrification in estuarine sediment microbial communities.

The Douro estuary, our case study, located on the northwestern Portuguese coast, is under constant anthropogenic pressure due to urban runoffs, sewage discharges, land reclamation and use of pesticides.^18^ The three main cities that border the estuary (*ca*. 700 000 inhabitants) contribute with three major wastewater treatment plants, with effluents being drained to the estuary. Douro River is an international commercial waterway and an increase in traffic, both commercial and touristic, has been registered in recent years.^19^ With this comes an increase in the amount of Cu NPs derived from antifouling paints that can be expected to be released into the river. In past decades Cu-based pesticides were used vastly in the Port wine region and ended up in the Douro watershed, causing concern over the amounts of Cu in the ecosystem.^20^ Nowadays, Cu is increasingly being applied as a nanopesticide,^5^ which in spite of the promise of a slower and more controlled release of the active ingredient, still needs to be reapplied to the vineyards to offset the rainfall wash-off just as traditional pesticides did, contributing a point source of Cu NPs in the Douro watershed.

## 2 Experimental

### 2.1 Study site and sample collection

Douro estuary is located on the north western Portuguese coast (Fig. 1). It is a mesotidal estuary subjected to North Atlantic meteorological and hydrodynamic conditions.^21^ The mouth of the estuary is narrow and two thirds of it is shielded by a sandy spit, creating an ecosystem of great biological interest. For that reason the “Nature Reserve of the Douro Estuary” was step up in 2007 with the aim of protecting the birds and the landscape of this salt marsh.

**Fig. 1.**
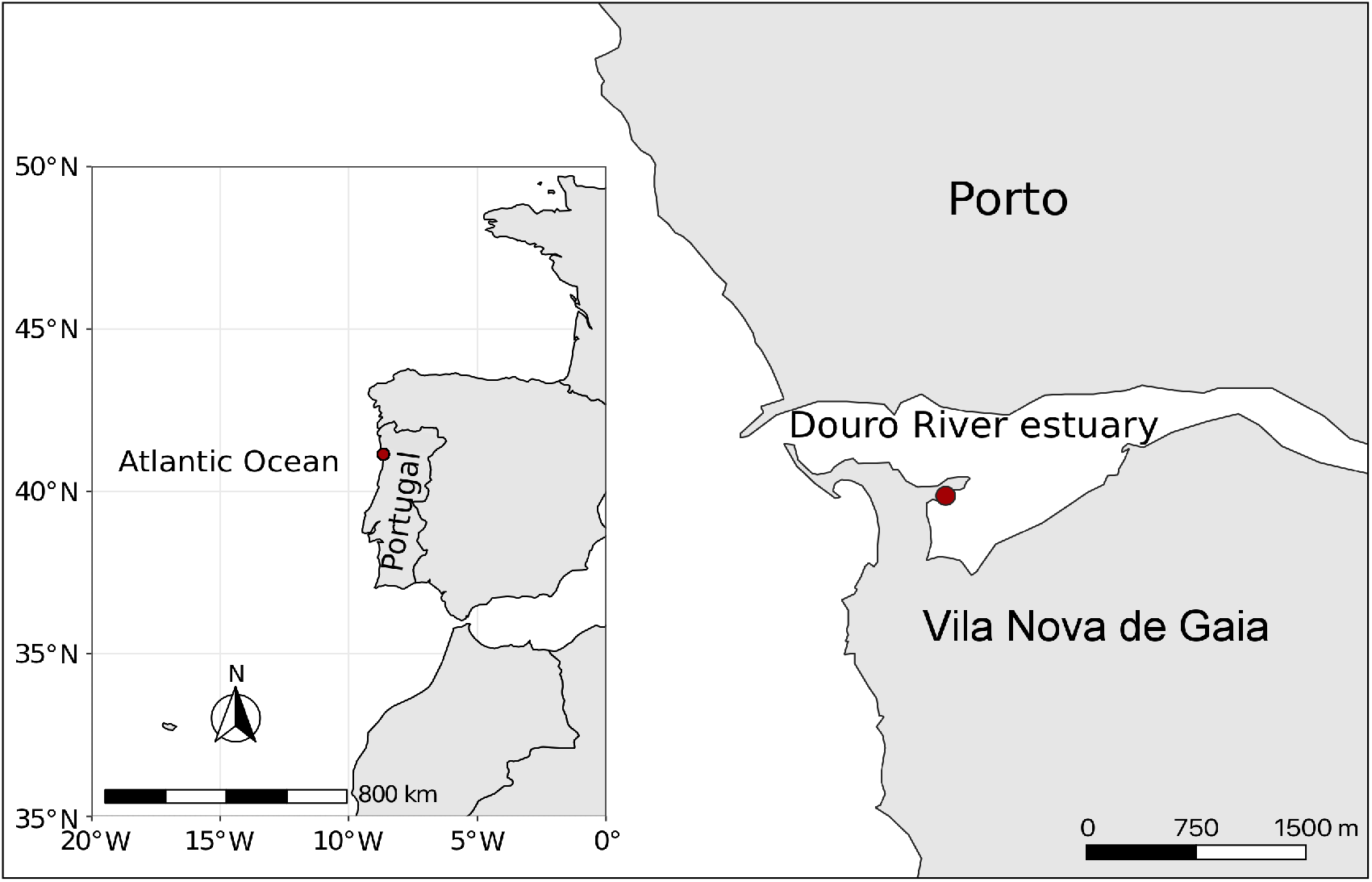
Study area and sampling site. Map of the Douro river estuary detailing the location of the sampling site (red dot) and inset map of continental Portugal and the Atlantic Ocean, where Douro disembogues.

Sample collection took place at the salt marsh (41°8’28.2588”N 8°39’48.2328”W), taking care to minimise disturbance to wildlife and habitats.^22^ This site was chosen as it had been characterized in previous studies and showed a sediment grain mostly composed of coarse sand (> 0.5 mm) and concentration of Cu < 4.5 μg g^-1^.^16^ The impact of Cu on denitrification had also been assessed^16^ as well as the impact on diversity, abundance and transcription of genes of the denitrification pathway,^17^ making this site very suitable to compare these data with data derived from Cu NPs exposure.

At low tide, intertidal sandy sediment (*ca*. 3 kg) were collected with an acrylic core (10 cm diameter) at 10 cm depth and homogenized in sterile Whirl-Paks. Overlying water (*ca*. 5 L) was collected and stored in acid-cleaned Nalgene bottles. Water temperature, dissolved O_2_, conductivity, salinity and pH, were measured *in situ* at the time of sampling (Table 1), with a multiparameter probe (pHenomenal^®^ MU 6100 H, VWR). Samples were transported to the laboratory refrigerated and in the dark.

**Table 1.**
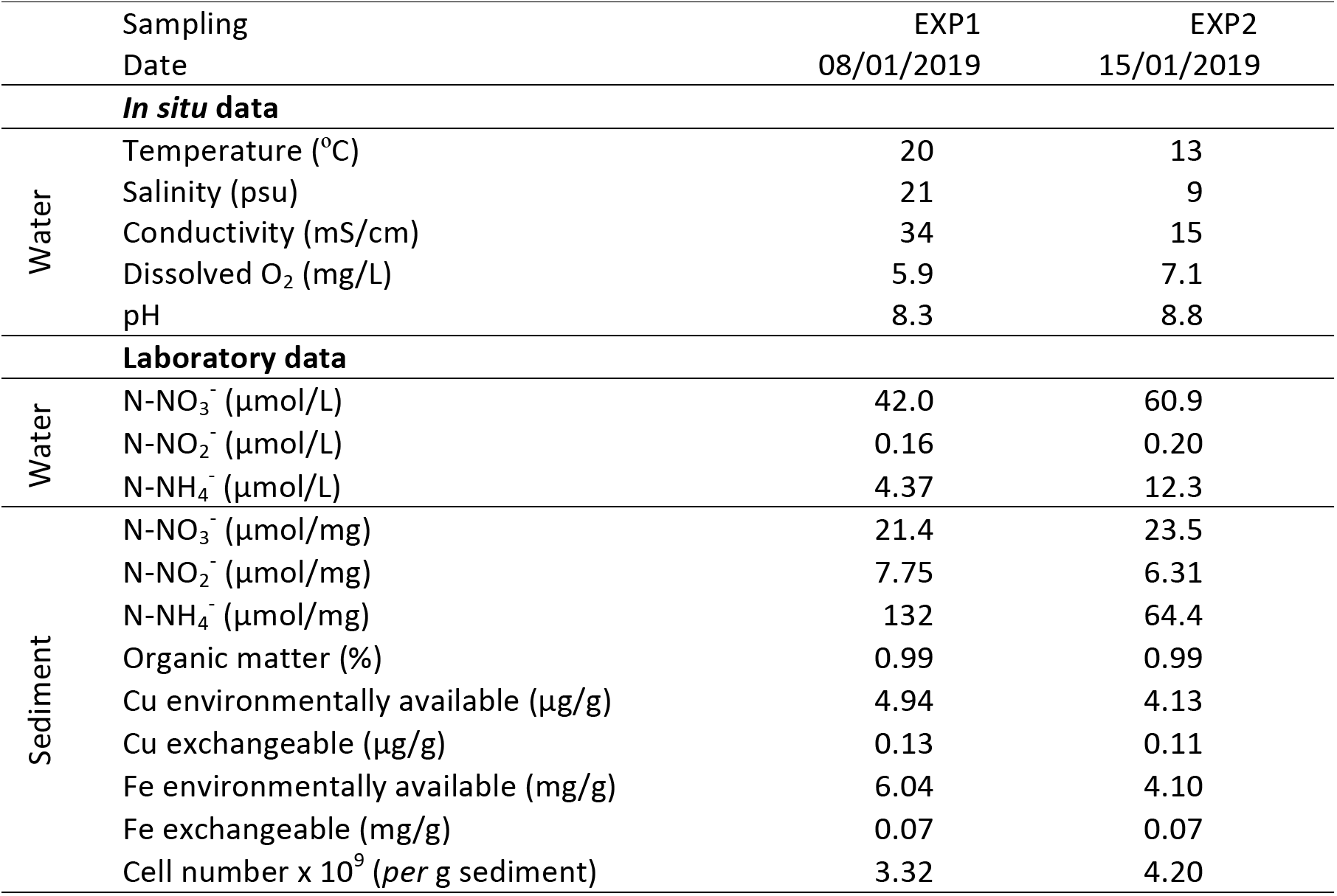
Water and sediment characteristics at the time of sampling.

### 2.2 Nanoparticle dispersions

Cu NPs were purchased from Sigma-Aldrich as a copper (II) oxide nanopowder, < 50 nm particle size (TEM) and a copper-zinc alloy nanopowder, < 150 nm particle size (SEM), 56-60 % Cu basis, 37-41 % Zn basis (reference numbers 544868 and 593583, respectively). To produce 100 mg L^-1^ stock dispersions 10 mg of Cu NPs were added to 1 mL of deionized water, sonicated for 30 min, vortexed for 30 s, diluted to 10 mL with filtered estuary water (0.22 μm cellulose membrane, Merck Millipore) containing 10 mg L^-1^ of chitosan and acetic acid (2 %), and again vortexed for 30 s.^12^ The obtained dispersions were characterized in the Instrumental Laboratory of the Department of Chemistry and Biochemistry, Faculty of Sciences, University of Porto. The hydrodynamic size (DLS) was 33 ± 12 nm and 119 ± 50 nm for Cu NPs < 50 nm and < 150 nm, respectively. Cu was analysed by flame atomic absorption spectroscopy (AAS) and the measured amount corresponded to 70 % and 40 % of the initially supplied, for Cu NPs < 50 nm and < 150 nm, respectively. Working Cu NPs dispersions were prepared daily by diluting the stock with filtered (0.22 μm) estuary water.

### 2.3 Microcosms

Microcosms were established in acid-cleaned glass serum bottles with 60 g of homogenized sediment and 60 ml of filtered estuary water amended with working Cu NPs dispersions to achieve final concentrations of 0.01, 0.1 or 1 μg g^-1^. Microcosms without Cu NPs addition were prepared in parallel and served as control. For Cu NPs < 50 nm microcosms were setup with overlaying water of two different salinities, 21 psu (EXP1) and 9 psu (EXP2), respectively, while for Cu NPs < 150 nm microcosms were setup only with overlaying water of 9 psu salinity. The microcosms were incubated for 6 days with constant agitation (150 rpm), in the dark. At 1, 4, 7, 24, 48 and 144 h one microcosm *per* concentration was used to collect water and sediment samples for analysis. At that time water physicochemical parameters were determined as described for the sampling.

Water and homogenized sediment aliquots were kept at −20 <u>°</u>C for analysis of inorganic N compounds, Cu and Fe concentration, organic matter amount and total bacteria enumeration. For analysis of inorganic N compounds sediments were dried at 105 °C until constant weigh^23^ prior to the analysis. Inorganic ammonium (N-NH_4_^+^), nitrite (N-NO_2_^-^) and nitrate (N-NO_3_^-^) were spectrophotometrically quantified (UV mini 1240 Shimadzu) on both sediment and overlaying water, in triplicate for each sample, as previously described^24^ and using freshly prepared standards.^25^ The limit of detection was 1, 0.1 and 2 μmol L^-1^ for N-NH_4_^+^, N-NO_2_^-^, and N-NO_3_^-^, respectively, and the relative standard deviation (RSD) between replicates varied between 3 and 7 %. Consumption or production rates of inorganic N compounds were calculated from the slopes of the linear regressions of concentration *vs* time. Fe and Cu concentrations in the sediment were analysed in the fraction termed “environmentally available” (digested at high pressure with nitric acid) and in the fraction termed “exchangeable” (room-temperature extracted with acetic acid) as previously described.^26^ Samples were analysed in triplicate by AAS, either with flame (Fe) or with electrothermal atomisation (Cu), at the Instrumental Laboratory of the Department of Chemistry and Biochemistry, Faculty of Sciences, University of Porto. Aqueous-matched standards were used for external calibrations. Quality control checks included analysing blanks containing no sediment alongside the samples. The limit of detection was 0.07 μg g^-1^ and 0.005 mg g^-1^ for Cu and Fe, respectively, and the RSD between replicates varied between 3 and 5 %. Organic matter (OM) content was determined by drying the sediments until constant weight, followed by ignition in a muffle furnace at 500 °C for 4 h and then reweighing. For total bacteria enumeration homogenized sediment (*ca*. 0.5 g) was treated with a saline solution (9 g L^-1^ NaCl) containing 12.5 % (v/v) of Tween 80 and fixed with 4 % (v/v) of formaldehyde. The slurries were stirred at 150 rpm for 15 min, followed by sonication for 20 - 30 s, at a low intensity. Subsamples of the slurries were then stained with 0.5 mg mL ^-1^ of DAPI (4’,6-diamidino-2-phenylindole) and incubated in the dark for 12 min. Samples were filtered onto black polycarbonate filters (0.2 μm Whatman Nuclepore) and washed with distilled water under gentle vacuum. All used solutions were 0.2 μm-filtered and when appropriate autoclaved. Membranes were set up in glass slides and cells were counted on an epifluorescence microscope (Olympus BX41 coupled to a U-CMAD3 camera adapter). A minimum of fifteen pictures were taken running through the entire slide (inverted S) and subsequent cell enumeration was performed with ImageJ.^27^

Sediment aliquots for nucleic acids extraction were kept in LifeGuard Soil Preservation Solution (Qiagen) at −80 °C. Total RNA and DNA were extracted from approximately 2 g of sediments with RNeasy PowerSoil Total RNA Kit and RNeasy PowerSoil DNA Elution Kit (Qiagen), respectively. RNA was reverse transcribed using the Maxima First Strand cDNA Synthesis Kit (Thermo Scientific). Kits were used according to the manufacturer’s instructions. Quantification of nucleic acids was performed with Qubit 3.0 using either ssDNA or dsDNA HS Assay Kit (Invitrogen), as appropriate.

### 2.4 Quantification of gene expression and abundance

An *in silico* analysis was carried out based on data from a diversity assessment (*16S rRNA* gene) previously performed at the Douro estuary,^28^ in order to choose the primers with the widest possible coverage (*Supplementary Information*, Table S1). Based on the results we selected the primers that target sequences affiliated with *nirK* in Cluster II (nirKC2F/nirKC2R) and *nirS* in Cluster I (nirSC1F/nirSC1R),^29^ and *nosZ* in Clade I (nosZ2F/nosZ2R),^30^ as being the most representative in our dataset. The *16S rRNA* gene was amplified with the bacterial primers 341F/534R and was used as reference gene.^31^

To determine gene expression *nirS* and *nosZ* were amplified from the cDNA samples by reverse transcription quantitative real-time PCR (RT-qPCR), and *nirK* was amplified from the DNA samples by qPCR, both performed in a StepOnePlus real-time PCR System (Applied Biosystems) using Power SYBR Green PCR Master Mix (ThermoFisher Scientific). A serial dilution of a template pooled from six to eight microcosms samples was always included in order to estimate the amplification efficiency of each gene with each sample.^32^ The slope and correlation coefficients of the standard curves and the PCR efficiency (E) were calculated with the built-in software of the equipment (software v2.3). Relative quantification of gene expression was performed by the 2^-ΔΔ*C*T^ method,^33^ assessing the change in the expression of *nirK, nirS* and *nosZ* relative to the control without Cu NPs addition, and using *16S rRNA* as reference, after ascertaining that its expression did not change over time. Differentially expressed genes were identified by taking into account the number of standard deviations by which a value was above or below the mean log2 fold change.^34^ This allowed defining a global fold-change difference and confidence and genes were considered differentially expressed when the log2 fold change had an absolute value higher than four. Absolute quantification to determine gene expression in numbers of copies was performed with a standard curve obtained from DNA extracted from bacteria *Roseobacter denitrificans* (DSMZ-German Collection of Microorganisms and Cell Cultures GmbH), after ascertaining it had the functional genes *nirS* and *nosZ*, and cloned using the TOPO-TA cloning kit with pCR 2.1-TOPO and One Shot TOP10 Chemically Competent *E. coli* (Invitrogen – Thermo Fisher), as described in the *Supplementary Information*. Results are reported in copy number *per* gram dry sediment, where the weight of the sediment used in DNA extraction was corrected for the water content. The change in abundance of *nirS* and *nosZ* was defined as the ratio of the log10 of gene copies to *16S rRNA* copies. Samples were run in duplicate, controls without template were always run in parallel with the samples, and the melting curves were always analysed to confirm that only target genes were quantified.^35^ Information regarding reaction mixtures and thermal programs is summarized in the *Supplementary Information*, Table S2, following the MIQE guidelines.^36^

### 2.5 Measurement of N2O production rates and determination of potential denitrification activity

N_2_O production rates were measured using the acetylene (C_2_H_2_) inhibition technique. Homogenized sediment (5 g) were weighted into 50-mL serum bottles and covered with 10 mL of 0.22 μm-filtered estuarine water, amended with KNO_3_ (to a final concentration of 50 μmol L^-1^) and with the respective Cu NPs dispersion to achieve final concentrations of 0.01, 0.1 or 1 μg g^-1^. Controls consisted of sediments without Cu NPs addition. Serum bottles were purged with N_2_ for 15 min and C_2_H_2_ was injected (15 % of the headspace) into the bottles. Each treatment was incubated with and without C_2_H_2_, for 4 h and 7 h, in duplicate, in the dark, at 150 rpm. The linearity of the process within this time frame had been confirmed before.^24^

Gas samples were collected from the serum bottles, after headspace equilibration via vigorous shaking, and immediately injected into a Varian gas chromatograph (CP-3800) equipped with an electron captor detector (ECD) held at 250 °C. Separation was achieved on two HayeSep D columns (0.9 and 1.8 m), kept at 80 °C, using a mixture of 5 % methane in argon as a carrier gas. Concentration was calculated using a certified standard with a concentration of 100 ppm (N_2_O in He). To account for N_2_O dissolved in solution Bunsen coefficients (T = 25 °C, P = 1 atm) were used to calculate the amount of gas dissolved in the liquid phase from the concentration in the gas phase.^37^ The amount of N_2_O produced was calculated by dividing by the wet weight of the sediment and N_2_O production rates were calculated from the slopes of the linear regressions of concentration *vs* time. Potential denitrification rates were calculated from the N_2_O produced with and without C_2_H_2_. Inorganic N compounds concentrations were determined for each sample, as described in 2.3, and used to calculate consumption or production rates from the slopes of the linear regressions of concentration *vs* time.

### 2.6 Data analysis

R v.3.5.1 was used for the analysis^38^ with Base R and package ‘tidyverse’ v.1.2.1.^39^ Plots were produced using R packages ‘ggplot2’ v.3.2.0,^40^ ‘cowplot’ v.1.0.0^41^ and ‘scales’ v.1.1.0^42^. Maps were obtained on Google Maps (https://maps.google.com) and Natural Earth (https://www.naturalearthdata.com) and created with QGIS v.3.6^43^ and R packages ‘ggplot2’, ‘sf’ v.0.8.1,^44^ and ‘ggspatial’ v.1.0.3.^45^ Kruskal-Wallis tests were performed to compare the response of a variable across the different microcosms, after ascertaining that the data did not follow a normal distribution (Shapiro-Wilk test); whenever a difference was found a Wilcoxon rank-sum test, using a Benjamini-Hochberg correction for multiple testing, was conducted to test for pairwise differences between microcosms. Pearson’s correlation coefficient was employed to assess linear responses of microcosms parameters over time and the slopes of those responses were compared with a *t*-test. Estimation plots were computed and generated with R package ‘dabestr’ v.0.2.3.^46^ To assess dissimilarities between different microcosms data were standardized, euclidean distances were calculated with function *vegdist* in R package ‘vegan’ v.2.5.5^47^ and represented in a principal coordinate analyses (PCoA).^48^ Differences between groups were assessed with a PERMANOVA^49^ based on 999 permutations and performed with the function *adonis2*, after testing for the homogeneity of group dispersions with the function *permutest.betadisper*, both in R package ‘vegan’. Data and code for the analysis and figures are available at https://github.com/msbaptista/Copper_nanoparticle_estuary

## 3 Results

### 3.1 Microcosms conditions over the time of exposure

In this work the estuary water never displayed salt wedge characteristics, *i.e*. salinity equal to or higher than 30 psu (Table 1). Douro estuary has been described as a salt wedge. However, for the lower estuary section where the sampling took place, the oceanic water intrusion has been shown to be absent from early December until late March.^21^ Accordingly, in these circumstances it was considered that when assessing the response of specific genes of the denitrification pathway, salinity would not be a factor of variability. To back this assumption microcosms exposed to Cu NPs < 50 nm set up with overlaying water of either 21 or 9 psu salinity were compared. The influence of salinity in the denitrification process has been considered as one of the most important environmental constraints of this N cycle pathway in estuaries.^50^ Salinity is also known to potentiate the mobility of Cu in soils and sediments,^51,52^ and to greatly influence the behaviour of Cu NPs.^8^

A PCoA showed that Cu concentration over time displayed the same behaviour for both salinities (Fig. 1S), and a PERMANOVA showed that differences in Cu concentration for both salinities were not significant (*F* = 0.420) whereas for time points a slightly significant difference could be seen (*F* = 1.571). Testing the data dispersion showed it was not homogeneous (*F* = 3.244), and thus the significant result for time points was interpreted as reflecting the dispersion of the data instead. To further investigate the effect of time, linear regressions of the physicochemical parameters in the microcosms were analysed. Only pH and N-NO_3_^-^ concentration in water increased in a time dependent manner (*p*-value <0.001). A *t*-test showed no differences in the increases in the two salinity regimes (*p*-value = 0.732 and 0.179, for pH and N-NO_3_^-^, respectively), showing the temporal dynamics to be similar regardless of the salinity value.

After ascertaining that salinity was neither influencing the microcosms response during the time-course experiment, neither the Cu concentration, the size and amount of Cu NPs, and number of bacterial cells in the sediment were the variables considered to assess the response of the microcosms throughout the exposure period. A PCoA showed that, for both Cu NPs < 50 nm and < 150 nm, samples within the first 24 h were grouped closer than after 48 or 144 h (Fig. 2A). In accordance, a PERMANOVA showed that differences in Cu concentration were not significant (*F* = 1.086) whereas differences in time and size were (*F* = 4.386 and 16.71, respectively). Since the dispersions of these groups were not significant (*F* = 0.828 and 0.595 for time and size, respectively) the results of the PERMANOVA were interpreted as reflecting differences in the microcosms exposed to different sizes of Cu NPs over time. Once again, regarding the physicochemical parameters, pH and N-NO_3_^-^ concentration in water increase over time (Table S3). Comparing these increases in microcosms exposed to Cu NPs < 50 nm and < 150 nm with a *t*-test showed that only N-NO_3_^-^ concentration in water had a different behaviour for both sizes (*p*-value = 7.40E-05), with a slightly higher increase in the microcosms exposed to Cu NPs < 50 nm.

**Fig. 2.**
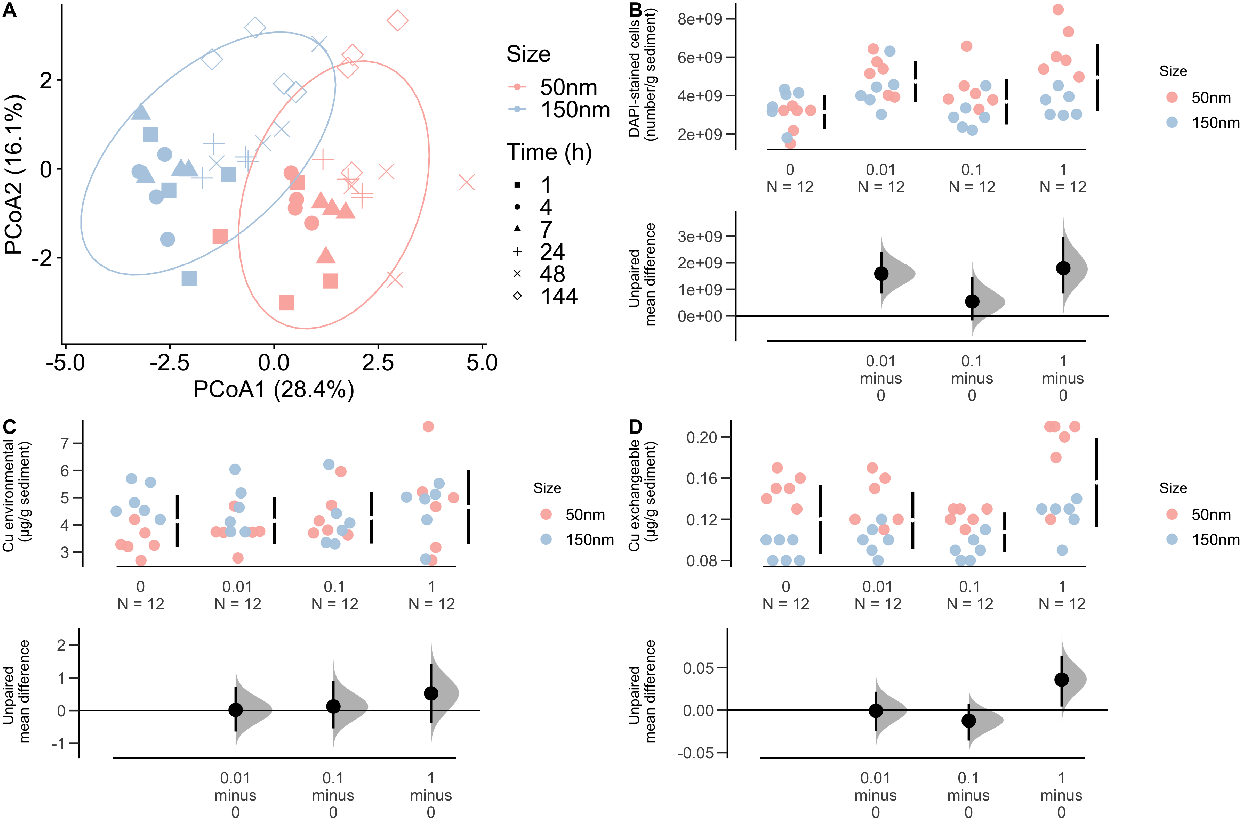
Copper availability and cell numbers in the microcosms exposed to Cu NPs. **A)** Principal component analysis, with ellipses overlaid at a 0.95 level, showing the separation between the microcosms exposed to Cu NPs < 50 nm and < 150 nm, particularly in the first 24 h of exposure; **B)** Number of bacteria (DAPI-stained cells) in the sediment; **C)** Cu concentration in the sediment environmentally available fraction and **D)** in the sediment exchangeable fraction. In the estimation plots (B-D) results are reported for microcosms exposed to Cu NPs 0, 0.01, 0.1 and 1 μg g^-1^. Gapped lines to the right of each concentration group are mean value ± standard deviation and the effect size, with 95 % confidence intervals, is plotted below.

Potential toxicity for prokaryotes resulting from the addition of Cu NPs to the sediment was not expected, given the range of concentrations tested and what is currently known regarding toxicity effects of Cu NPs towards the aquatic biota.^2^ In line with this assumption no toxicity of Cu NPs, evaluated by a decrease in number of bacteria, could be seen. Quite the opposite, bacterial cell numbers were slightly higher (Kruskal-Wallis test, *p*-value = 0.008) in microcosms exposed to Cu NPs < 50 nm than in microcosms not exposed to Cu NPs, with the estimation plots showing higher average cell number for microcosms exposed to 0.01 and 1 μg g^-1^ Cu NPs < 50 nm (Fig. 2B).

Cu concentration in the sediments used to establish the microcosms was 4.5 ± 0.7 μg g^-1^ (n = 6), similar to previous reports at the same sampling location,^16^ which shows that Cu concentration has remained fairly constant throughout the years. Taking into account that environmental levels of Cu were *ca*. 4 times higher than the highest ones tested in this work, and considering their variation (± 0.7), we did not expect to be able to detected differences in microcosms upon Cu NPs addition and measurement by AAS. In fact, Cu environmentally available fraction showed no differences between microcosms (Fig. 2C). Cu exchangeable fraction, however, seemed to increase slightly in microcosms exposed to 1 μg g^-1^ Cu NPs < 50 nm (Fig. 2D), the estimation plot showing the average value in these microcosms to be higher than the average value in microcosms not exposed to Cu NPs, hinting at a higher Cu bioavailability. However, for the Cu exchangeable fraction it is also possible to see a higher variability in the microcosms not exposed to Cu NPs. Cu exchangeable fractions previously reported for the sediments of Douro estuary and Douro tributaries showed great variation according, for instance, to the occurrence of algae blooms^26^ or plant colonization of the sediment.^53^ Since this extraction procedure assesses the metal fraction more weakly bound to the sediment it is more prone to variation, as weakly bound metals are released more stochastically. Fe environmentally available and exchangeable fractions in sediments did not show differences between microcosms. Cu environmentally available fractions in sediments are often normalised to the Fe fractions, since this element, particularly when employing total-recoverable digestions as we do in this work, can be appropriately used as a conservative tracer, and, moreover, this ratio can be a more useful tool in predicting disturbances in sediment biota communities than elemental concentrations alone.^20^ Normalised Cu concentrations did not vary in this work (Fig. 2S) reflecting the even impact Cu NPs addition to the microcosms.

Taken together these data indicate that the microcosms exposed for up to 6 days maintained similar physicochemical conditions in water and sediment exposed over time. A clear distinction between microcosms exposed to different Cu NPs sizes could be seen. For microcosms exposed to Cu NPs < 50 nm the number of bacterial cells and the Cu exchangeable fraction in the sediment increased in a concentration dependent manner, which for microcosms exposed to Cu NPs < 150 nm could not be seen.

### 3.2 Gene expression response to Cu NPs of different sizes

Further strengthening the differences between microcosms exposed to different Cu NPs sizes, a distinct pattern of gene expression could be seen. For microcosms exposed to Cu NPs < 50 nm the expression of *nirS* and *nosZ* increased in the first 24 h, whereas for microcosms exposed to Cu NPs < 150 nm no substantial change in gene expression could be seen (Fig. 3).

**Fig. 3.**
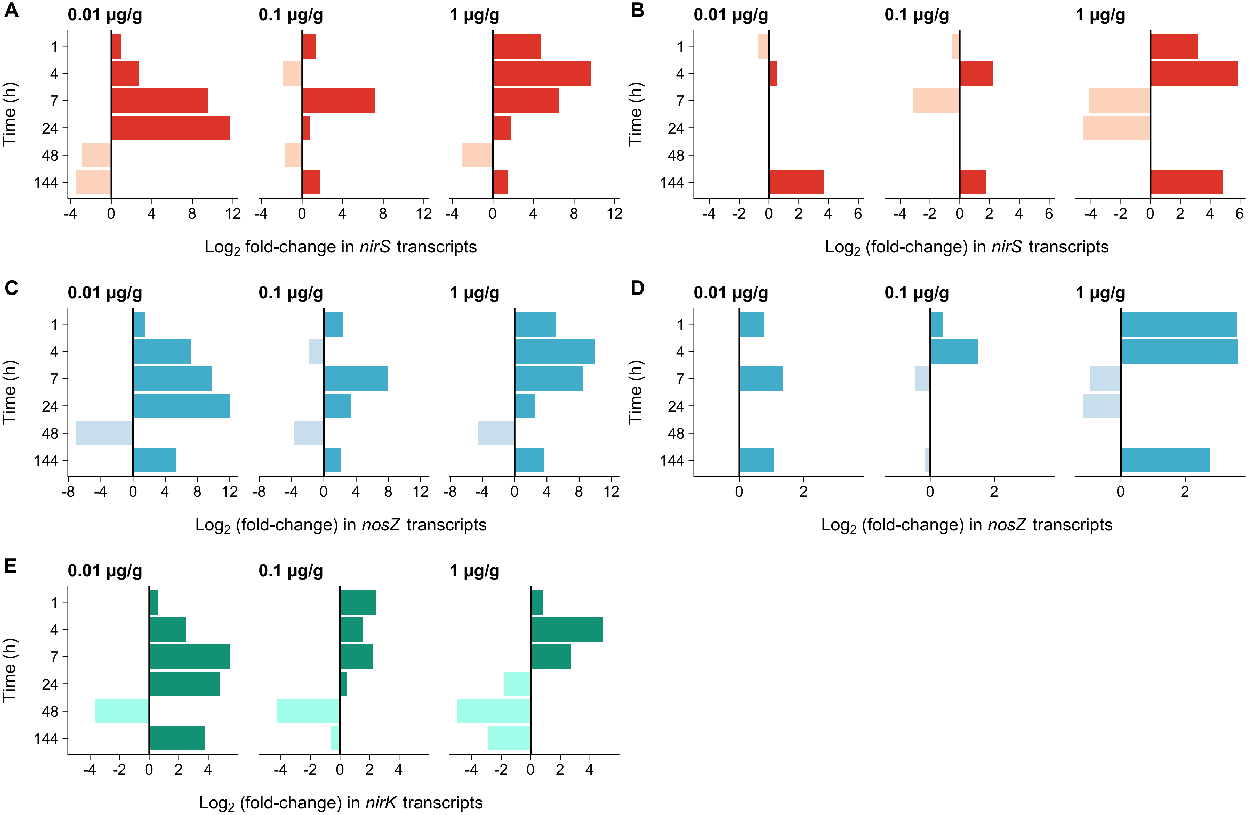
Changes in gene expression in microcosms exposed to Cu NPs. Relative quantification was determined by fold changes in expression of **A)** *nirS* transcripts in microcosms exposed to Cu NPs < 50 nm; **B)** *nirS* transcripts in microcosms exposed to Cu NPs < 150 nm; **C)** *nosZ* transcripts in microcosms exposed to Cu NPs < 50 nm; **D)** *nosZ* transcripts in microcosms exposed to Cu NPs < 150 nm; **E)** *nirK* transcripts in microcosms exposed to Cu NPs < 50 nm. Fold change in gene expression is the average of two replicates; RSD between replicates varied between 3 and 17 %.

For microcosms exposed to 0.01 μg g^-1^ Cu NPs < 50 nm substantial increases in gene transcripts could be seen after 4 h (*nosZ*) or 7 h (*nirS*) and continuing to increase up to 24 h, when the highest expression was obtained. For 0.1 μg g^-1^ a peak in expression could be seen after 7 h of exposure, and for 1 μg g^-1^ increases in expression could be seen immediately after 1 h of exposure and continuing to increase up to 4 h, time after which the expression decreased (Fig. 3A-C). *nirK* expression in microcosms exposed to Cu NPs < 50 nm followed roughly the same pattern seen for *nirS* and *nosZ*, but the smaller log2 fold change in gene transcripts was interpreted as not representing significant differences compared to the control (Fig. 3E). The same smaller log2 fold change in gene expression could be seen for all the microcosms exposed to Cu NPs < 150 nm, which again was interpreted as not representing significant differences to the control (Fig. 3B-D). Given that the gene expression analysis was performed in a sequential manner, starting with microcosms exposed to Cu NPs < 50 nm, it was considered not worth it to analyse *nirK* transcripts in microcosms exposed to Cu NPs < 150 nm, as overall analysis of the data suggested that those would not be different from the control. After 48 h of exposure no changes could be seen in gene expression for any microcosm exposed to Cu NPs, a situation that was still maintained by the end of the of exposure, after 6 days, corroborating the results seen with the PCoA analysis (Fig. 2A) that samples obtained in the first 24 h were separated from the subsequent ones.

Gene expression showed a strong dependence of time and Cu NPs < 50 nm amount. The earliest increases in gene expression occurred for the highest Cu NPs additions, for which a peak was achieved after 4 h of exposure. Conversely, the lowest Cu NPs additions only showed increases in gene expression after 4 h or 7 h of exposure, but that continued to increase until 24 h achieving an overall higher gene expression. To check if the addition of Cu was reflected at the gene level, we investigated *nirS* and *nosZ* copy numbers for the microcosms exposed to Cu NPs < 50 nm. As can be seen in Fig. 4(A-B), the increase in gene copy numbers followed the same temporal and concentration trends described for the gene transcripts, with 0.01 μg g^-1^ showing a peak after 24 h, 0.1 μg g^-1^ showing a peak after 7 h and 1 μg g^-1^ showing a peak after 4 h of exposure, and with the highest gene copy number obtained for 0.01 μg g^-1^ after 24 h. When normalizing *nirS* and *nosZ* by *16S rRNA* copy number, to account for fluctuations in the number of cells, it was possible to see an increase in the number of *nirS* and *nosZ* genes in microcosms exposed to increasing concentrations of Cu NPs < 50 nm (Fig. 4C-D). However, a higher variability of copy numbers could also be seen for the microcosms exposed, highlighting the diffused contribution of Cu NPs to the observed changes.

**Fig. 4.**
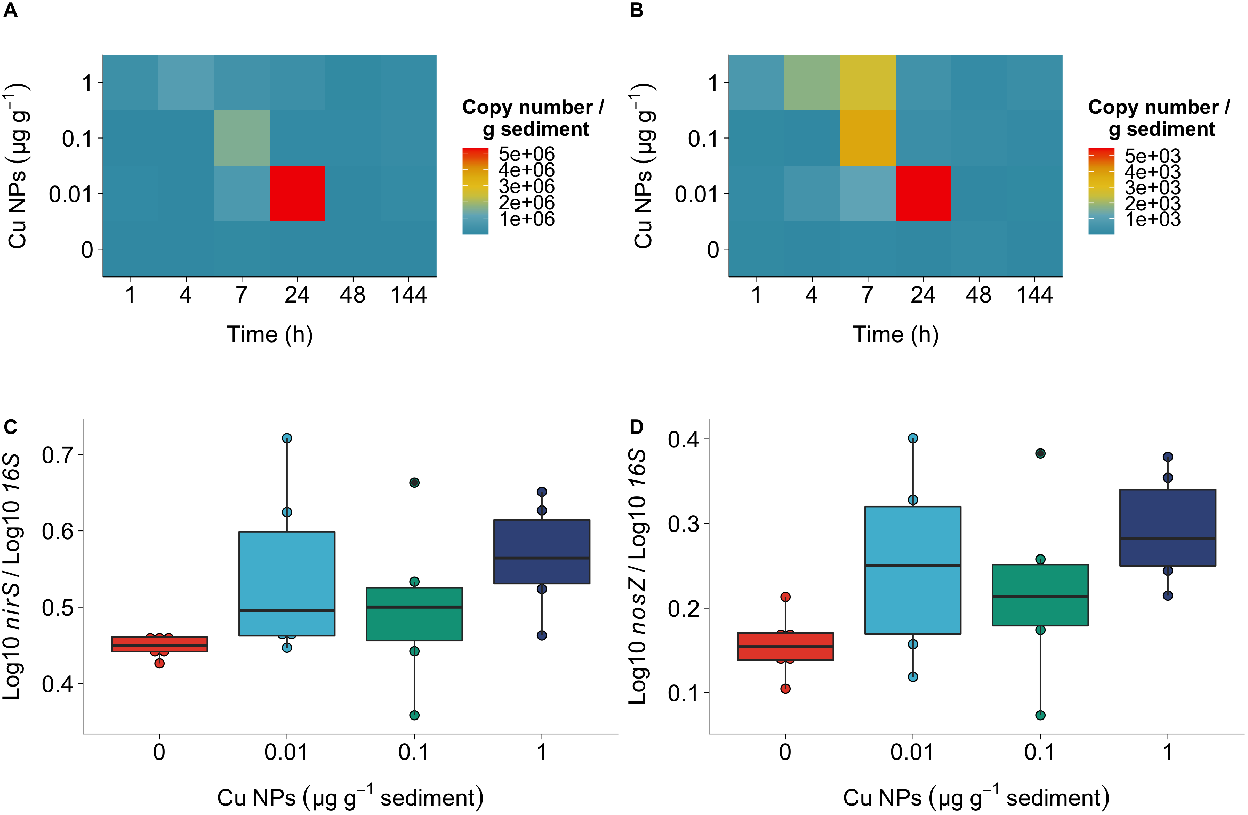
Abundance of denitrification genes in microcosms exposed to Cu NPs < 50 nm. Absolute quantification of gene copy number per gram dry sediment is shown during the time-course experiment as a heatmap for **A)** *nirS* and **B)** *nosZ;* and shown for the different concentrations tested as a boxplot for **C)** *nirS* and **D)** *nosZ* with gene copy number per gram dry sediment log10 transformed, and *nirS* and *nosZ* values normalized by the *16S rRNA* gene.

Studies comparing the abundance of *nir* and *nosZ* in various environments (*eg*. soils, sediments) have shown that the number of *nir* genes can exceed that of *nosZ* by orders of magnitude,^54^ as well as that *nirS* can be consistently more abundant than *nirK* in estuaries.^55^ These same findings could be seen for this work, showing these to be the conditions of Douro estuary and showing that these were captured on the microcosms setup. On microcosms not exposed to Cu NPs *nirS* abundance showed a correlation to bacterial *16S rRNA* abundance (Pearson’s *r* = 0.511, *p*-value = 0.110), whereas *nosZ* did not (Pearson’s *r*=0.012, *p*-value = 0.836); moreover *nirS* and *nosZ* were also not correlated with each other (Pearson’s *r*=0.221, *p*-value = 0.347). Conversely, for the microcosms exposed to Cu NPs, *nirS* and *nosZ* abundances were positively correlated for all three concentrations tested (Fig. 3S), suggesting a similar response from the denitrifier community. Known sequenced genomes show that most microorganisms carry one copy of either *nirK* or *nirS*.^56^ However, from the genomes that possess *nir* genes approximately two thirds is currently known to harbour *nosZ* as well.^57^ Therefore, the correlations obtained with this work hint at the possible presence of both genes in the better part of the denitrifier community of Douro estuary.

### 3.3 Potential denitrification in sediments exposed to Cu NPs

The potential denitrification rate (N_2_ plus N_2_O production) was determined for the microcosms for which a change in gene expression had been seen, namely those exposed to Cu NPs < 50 nm. The results showed an overall average of 30 % higher in sediments exposed to Cu NPs (Fig. 5A), increasing by 22 %, 24 % and 42 %, for 0.01, 0.1 and 1 μg g^-1^, when compared to the non-exposed control. The consumption of N-NO_2_^-^ was highest for sediments exposed to 1 μg g^-1^ (Fig. 5B), which showed the highest potential denitrification rate as well. For the remaining concentrations, higher productions of N_2_ plus N_2_O were associated with slightly higher consumptions of N-NO_2_^-^ for 0.01 μg g^-1^, however, for 0.1 μg g^-1^ the same trend could not be seen.

**Fig. 5.**
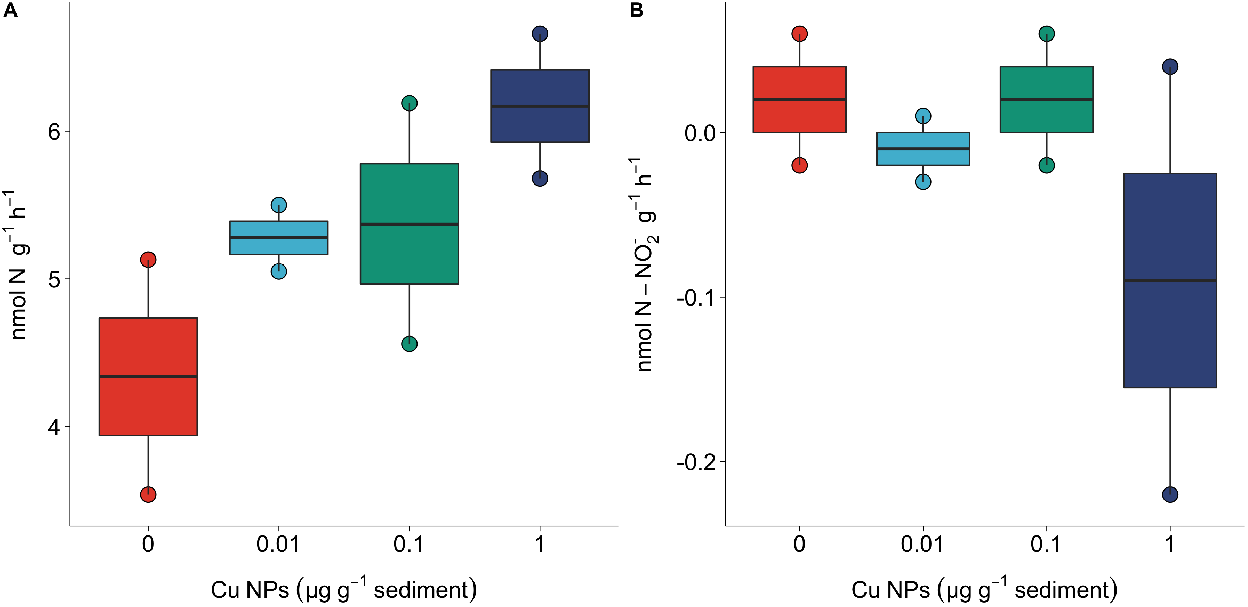
Potential denitrification and N-NO_2_^-^ consumption on the sediments exposed to Cu NPs < 50 nm. **A)** Potential denitrification rates were calculated from the difference in N_2_O produced with and without C_2_H_2_, over time; and **B)** N-NO_2_^-^ consumption rates were determine by the slopes of the linear regressions of concentration over time.

Gene expression up until 7 h was swifter under 1 μg g^-1^ exposure (Fig. 3A & C), and these results are reflected in the potential denitrification and N-NO_2_^-^ consumption rates that were seen. Though the highest gene expression seen in this study was for 0.01 μg g^-1^ after 24 h, incubations to determine potential denitrification rates did not continue up to this hour due to the fact that these would fall outside the linearity of the employed method.^24^ Linear accumulation of N_2_O is used as evidence for the validity of the results, as the acetylene inhibition technique is known to have time dependent bias that contribute to underestimating the denitrification rates.^37^ Specifically the inhibition of N_2_O-reductases by acetylene is incomplete, chiefly when soils are wet, are low in N-NO_3_^-^ or have a high C: N-NO_3_^-^ ratio. Also, acetylene has been reported to inhibit key enzymes involved in nitrification (ammonium monooxygenase), thus affecting denitrification rates when the amount of N-NO_2_^-^ or N-NO_3_^-^ available for this pathway is compromised.^58^ To overcome these issues additions N-NO_3_^-^ to are often performed when estimating the denitrifying potential of the sediments^37^ and, in this work, doing that resulted in the pool of N-NO_3_^-^ remaining unchanged up to 7 h.

Other studies^59^ at Douro estuary have shown that the variability in N-NO_3_^-^ concentrations modulates the magnitude of the denitrifying process, with potential denitrification rates ranging from 0.4 to 38 nmol N g^-1^ h^-1^ and varying greatly according to the different substrates studied, time of the day, or season.^24^ Rates measured with the isotope pairing technique, a more powerful tool to investigate rates of microbial mediated nitrogen processes^60^ has shown denitrification at Douro estuary to vary between 4 - 8 nmol N g^-1^ h^-1^,^61^ encompassing the ranges obtained in this study, and contributing to corroborate that, in spite of its simplicity, the acetylene inhibition method is a useful tool for relative comparisons of denitrification rates.

## 4 Discussion

In this work we addressed two noteworthy issues regarding the effect of Cu NPs upon estuarine microbial communities that have often been overlooked, by looking into the effects of environmentally-relevant concentrations and by assessing the impact of Cu NPs not only on the microbial communities but on the ecosystem services they provide, namely the service of removing fixed nitrogen, which is critical in eutrophic ecosystems.

Previous works have shown that the size of metallic NPs could better predict its toxicity than the metal content^62^ and in accordance our data shows that microbial communities in the estuarine sediment responded to additions of Cu NPs < 50 nm, but not to additions of Cu NPs < 150 nm. The characterization of the NPs in the stock dispersion showed sizes of *ca*. 30 and 120 nm, for Cu NPs < 50 nm and < 150 nm, respectively. Therefore, Cu NPs < 150 nm were still larger than 100 nm, a size that has often been described as the upper limit for a material to be considered a NP and, thus, to behave as such.^63^ The Cu NPs < 150 nm used in this work were an alloy with *ca*. 60 % Cu, which was reflected in the amount of Cu measured in the NPs dispersions (40 % of the nominal concentration *vs* 70 % for Cu NPs < 50 nm). Despite this, the estimated actual tested concentrations for both NPs sizes were of the same magnitude, and ranged from 7 x 10^-3^ −10^-1^ μg g^-1^ to 4 x 10^-3^ −10^-1^ μg g^-1^, for Cu NPs < 50 nm and < 150 nm, respectively. Cu NPs have been predicted to occur in the environment, specifically in freshwater and seawater systems, in ranges of μg *per* kg, with the lowest Cu amount tested in this work corresponding to the upper predicted limits.^2,64^ At these values no toxicity towards the estuarine sediment bacterial community could be seen. Notwithstanding the other tested concentrations being above the upper limit of predicted environmental concentrations, they also did not show a toxicity effect. Other studies have shown no toxic effects on bacteria exposed up to 10 mg L^-1^ Cu NPs, both on freshwater and seawater,^9,65^ indicating a high tolerance to Cu in the NP form. The nature of the antimicrobial activity of metal-containing NPs is still under debate, particularly whether toxicity is mostly caused by the release of metal ions, by the disruption of the outer cell membranes integrity, or by the NP penetrating biological systems and contributing to cytotoxicity and genotoxicity, and the formation of reactive oxygen species. In a microcosms exposed to 50 nm Cu NPs (pH = 7.8, comparable to this work’s pH = 8) the dissolved Cu was shown to increase in a time dependent manner, with highest dissolution rates in the first 24 h, but consistently more than 50 % of the Cu remained undissolved and as a NP, for up to five days of exposure.^65^ These data suggest that in this work the Cu NPs remained predominantly as a NP in the microcosms and that the effects seen on gene expression were caused by their intrinsic properties rather than by the release of metal ions and corresponding dissolution.

The better part of the studies pertaining the impact of Cu NPs (and other metallic NPs) on biota have focused on establishing effect concentrations in order to predict exposure reference doses that could inform risk assessment frameworks and regulations.^7^ Instead, in this work we enquired what possible implications the existing amounts of Cu NPs in the ecosystem could have towards the biota. Implications for the estuarine microbial communities were considered based on the fact that genes of the denitrification pathway can be regulated in response to the extracellular Cu.^15^ Copper is a necessary co-factor in key enzymes of the denitrification process, namely the nitrous oxide reductase (NosZ) and the nitrite reductase (NirK). NosZ shows very similar properties when purified from a wide range of bacteria of diverse metabolic groups.^66^ For NirK, however, three different classes have been suggested based on differences on structure, but several known NirK have structures that do not fit any of the current classes.^55^ The co-occurrence of *nosZ* with *nirS* in the aquatic environment is more frequent than the co-occurrence with *nirK*,^67^ and an analysis of the Douro estuary microbial community showed that for Douro estuary this was the case (*Supplementary Information*, Table S1). Moroever, the concomitant increase in gene expression and copy number of both *nirS* and *nosZ* suggested that the community was composed by microorganisms that harboured both genes. Therefore, considering *nirS* and *nosZ* co-occurrence within the same microorganism, the impact of Cu NPs could have meant more available Cu to drive the expression of *nosZ*, which in turn would lead to the expression of *nirS*, since its expression would be necessary for the production of the substrate on which the nitrous oxide reductase would act upon. The analysis of the Douro estuary microbial community also showed that *nirS* could potentially be present in genera accounting for 1.3 % of the overall relative abundance whereas *nirK* genera could potentially account for 0.2 % of the overall relative abundance, enabling the assumption that the microcosms community was mostly comprised of *nirS*-containing organisms. The corresponding impact of Cu NPs on the change of *nirK* expression might not have been captured for the microcosms community due to the reduced number of microorganisms within the denitrifying community that harboured this gene. It could also have happen that under the chosen exposure conditions the expression of *nirK* was not enhanced due to lack of impact from the Cu NPs.

In this work we have showed that ecosystem services could be impacted by Cu NPs by ascertaining that both an increase in the expression of genes of the denitrification pathway and in N_2_O production were seen, after exposure to NPs. This relationship, instrumental to improve the prediction of rates of denitrification, cannot always be shown. Other works have reported that an increase in the denitrifying activity is not always associated with an increase in transcript or gene abundance.^68–70^ One of the reasons for this might be the modular character of the denitrification pathway, in that an organism does not always possesses the set of enzymes necessary to carry out the entire process. At Douro estuary Cu amendments of 4 μg g^-1^ (equal to the environmental measured levels) led to a reduction of total N removal via denitrification, at the same time that the release of N_2_O was enhanced. Furthermore, this amendment yielded a decrease in the abundance of *nirK, nirS* and *nosZ* transcripts.^16,17^ Cultures of the representative denitrifying bacteria *Pseudomonas stutzeri* have shown that decreasing the Cu concentration resulted in the accumulation of N_2_O, compared to Cu-replete cultures.^71^ However, the highest concentrations of N_2_O were not produced in Cu-deficient cultures but at Cu levels for which the lowest proportion of N_2_O to N_2_ levels were generated.^72^ Increasing the Cu concentration also increased the expression of *nirS* and *nosZ*, up until a concentration for which the transcription was inhibited, suggesting a differentiated production of N_2_O by *P. stutzeri* at lower or higher levels of Cu.^72^ The results obtained in this work show Cu NPs at the tested concentrations increasing the expression of genes of the denitrification pathway and contributing to the overall removal of N from the Douro estuary as N_2_, since the tested concentrations of Cu NPs are not toxic and, therefore, do not inhibit *nosZ* expression.

The ecological implications of these findings concern the fact that N_2_O is a potent greenhouse gas, and also the one who contributes the most to the destruction of the ozone layer,^73^ and the fact that in estuarine environments Cu concentration gradients can often be found. The Douro estuary, similarly to many others, harbours sediments displaying concentrations of Cu that are much higher than any up-to-date or predicted input of Cu NPs. However, as this work showed, inputs of metallic NPs into the environment should not be disregarded based on the fact that they constitute only a small fraction of the total metal content of the ecosystem. A comprehensive review of NPs in soils found reports that these could lead to reduced diversity and function of soil microorganisms, or could have no measurable effect, or could have positive effects on soil microbial communities and functioning, leading to the conclusion that, above all, NP-microbe interactions are context-driven.^74^ The best way to address these inconsistencies is by ensuring that experiments are conducted under more realistic environmental conditions, which was attempted with this work. Limitations of this work include the fact that it did not enquire about Cu NPs internalization, or cell wall binding, in prokaryotic cells. In the future, this understanding of the interactions between microorganisms and Cu NPs, will enable disentangling the effects of Cu as NP or as ion, and will be invaluable to establish causality between exposure to Cu NPs and increased gene expression and N_2_O emissions.

## 5 Conclusion

Nanotechnology, operating at a scale much smaller than any other, has shown how much it can impact living beings *in vitro*; but *in vivo* studies are only just starting to grasp the many implications that nNPs can pose towards organisms. The macroscale at which we measure the better part of the processes we aim at studying might be inadequate to assess the microscale at which the prokaryotic communities thrive and change, and the nanoscale at which NPsoperate. In this work we have began to find a way of interconnecting these findings by showing that Cu NPs, at concentrations that when assessed by classic toxicity studies do not exhibit either toxic or hormetic effects, can actually have an effect on biological pathways and impact important steps of the N cycle. Future studies on the long term impact of continuous inputs of metallic NPs on estuarine environments will enable the understanding of the possible multiple outcomes for microbial ecosystem services.

## Conflicts of interest

There are no conflicts to declare.

## Acknowledgements

This work was funded by national funds through Fundação para a Ciência e a Tecnologia, Portugal (FCT) with the projects UIDB/04423/2020, UIDP/04423/2020, and PTDC/BIA-MIC/30131/2017 (NANOSED) also funded by the “Norte Portugal Regional Operational Programme” (Norte2020), under the PORTUGAL 2020 partnership agreement, through the European Regional Development Fund (ERDF). We would like to thank Eng. Paulo Faria for the opportunity to sample within the Nature Reserve of the Douro Estuary.

## Supplementary Information

### Supplementary Experimental

#### Selection of primers targeting nitrite and nitrous oxide reductase genes

Both nitrite (*nir*) and nitrous oxide (*nosZ*) reductase genes have been shown to exhibit phylogenetic distinct Clades and to harbour taxonomically diverse microorganisms.^1–4^ This has posed challenges to the design of primers that are both unambiguous and comprehensive of the denitrifier diversity. Common primers for nitrite reductase *nirK* gene are F1aCu/R3Cu^5^ and nirK2F/nirK5R;^6^ for *nirS* gene are cd3aF/R3cd^7,8^ and nirS2F/nirS4R.^6^ At the time these were designed the sequences available were mostly from Proteobacteria strains, which narrowed the diversity of these genes to this taxon. Nowadays they are still commonly used but advances in culture-independent methods to investigate the microbial community composition allowed the identification of *nir* genes across diverse taxonomic groups. A recent revision grouped *nir* genes into four Clusters for *nirK* gene and three Clusters for *nirS*.^2^ The previously described primers only target *nirK* and *nirS* in Cluster I, and to address this issue the authors have designed specific primers for each Cluster.^2^

The *nosZ* gene primers nosZ2F/nosZ2R^9^ have been designed to be as universal as possible, and in spite of its potential bias against gram-positive bacteria they have been widely used ever since. More recently *nosZ* have been grouped into two different Clades. Clade I consisted mostly in Proteobacteria sequences following the species phylogeny as previously observed, and Clade II consisted of sequences found among a diverse range of bacterial and archaeal phyla.^4^ Moreover, *nosZ* in Clade II has since been recognized as one comprising atypical nitrous oxide reducers found in various ecosystems.^10^

In order to choose the primers with the widest coverage for our study, we carried out an *in silico* analysis based on a microbial diversity assessment (*16S rRNA* gene) previously performed at the same sampling site of Douro estuary.^11^ Prokaryotic OTUs were categorized based on the presence of the functional genes within genomes that coded for proteins in the denitrification pathway, as compiled and reported before for *nir*^2^ and *nosZ*.^4^

The analysis showed bacterial OTUs predominantly affiliated with genera showing *nosZ* in Clade I (Table S1). For this reason the primer pair nosZ2F/nosZ2R was chosen for this study, based on the intuition that the primer set yielding the greater diversity would be the most suitable one.^9^ Denitrifying OTUs were affiliated with Bacteria genera showing *nirK* in Cluster I and II, however, for *nirS* only OTUs affiliated with Bacteria genera showing *nirS* in Cluster I could be predicted (Table S1). As expected, we found that OTUs affiliated with genera that could potentially harbour *nirS* were much more abundant than OTUs affiliated with genera that could harbour *nirK* (relative abundances of 1.3 % and 0.2 %, respectively).^12,13^ The most commonly used primers were designed for *nirS* and *nirK* in Cluster I because for several environments they are the most ubiquitous and abundant. This was the case for *nirS*, and preliminary tests in our samples showed that using primers nirSC1F/nirSC1R for *nirS* in Cluster I resulted in amplification (assessed by loading the PCR product in an agarose gel). Therefore, these primers were chosen for this study. Contrary to what was expected, at Douro estuary *nirK* in Cluster II was more abundant than in Cluster I. To confirm this assumption we tried to amplified our samples using primers nirKC1F/nirKC1R and nirKC2F/nirKC2R and *nirK* in Cluster II was the only for which we could see amplification (assessed by loading the PCR product in an agarose gel). Therefore, we chose primers nirKC2F/nirKC2R for this study.

For Archaea, *Nitrosopumilus* sp., a genus potentially containing *nirK* homologous genes,^14^ was found at Douro estuary. However, Archaea represented only 0.7 % of the OTUs at our sampling site. Specific primers have been designed for targeting the archaeal *nirK*, and although its transcription potential has been shown, no significant increase in transcript copy number was observed with increased denitrifying activity.^14^ For these reasons we chose not to include these primers in the present study.

#### Positive controls for the genes of interest

Bacteria *Roseobacter denitrificans* (obtained from the DSMZ-German Collection of Microorganisms and Cell Cultures GmbH) was used as positive control for *nirS* in Cluster I and for *nosZ*, and *Fulvivirga imtechensis* AK7 (obtained from the Japan Collection of Microorganisms) was used as positive control for *nirK* in Cluster II. These strains were chosen after an *in silico* PCR was performed by retrieving the sequences of the genes of interest annotated in UniProtKB (https://www.uniprot.org/) for these strains, and uploading it in Serial Cloner 2.6 (http://serialbasics.free.fr/Serial_Cloner.html) to align with the primers. Both strains were grown in marine agar medium (3 %) for 2 - 3 days, at room temperature. DNA was extracted with the E.Z.N.A. Bacterial DNA Kit (Omega Bio-tek) and the genes were amplified in a PCR reaction on a Veriti Thermal Cycler (Applied Biosystems) performed with 12.5 μL of DreamTaq PCR Master Mix (Thermo Scientific), with primers at a concentration of 1 μmol L^-1^ and 25 - 50 ng of DNA template, in a final volume of 25 μL. The primers are described in Table S2 and the thermal cycler conditions were as described in previous works for *nirK*,^2^ *nirS*,^15^ *nosZ*.^9^ PCR products of duplicate reactions were pooled together and the bands with the expected size - 457 bp for *nirK*, 425 bp for *nirS*, 267 bp for *nosZ* - were excised from an agarose gel (1.5 %), under UV light, with the QlAquick Gel Extraction Kit (QIAGEN). The excised bands were sequenced with Sanger sequencing technology at STAB VIDA (Porto, Portugal). The *nirS* gene was sequenced only in the forward direction (due to low sample volume) and the remaining genes were sequencing in both forward and reverse directions. Forward and reverse sequences were assembled with the CAP3 software,^16^ embedded into the Unipro UGENE software (Linux 64-bit version 1.32).^17^ The consensus sequences were then confirmed by blast searches against UniprotKB (accessed on 24/11/2019) using the blastx algorithm (http://www.ncbi.nlm.nih.gov).^18^

#### Absolute and relative quantification of gene expression levels

Reverse transcription quantitative real-time PCR (RT-qPCR) was performed using 6 ng of cDNA for *16S rRNA, nirS* and *nosZ* in a StepOnePlus real-time PCR System (Applied Biosystems) using Power SYBR Green PCR Master Mix (ThermoFisher Scientific). Relative quantification of gene expression levels was obtained by serial dilution of cDNA pooled from eight microcosms samples which was used as standard to allow calculating the slope and correlation coefficients of the standard curve and the PCR efficiency (Table S2). For *nirK* the same conditions resulted in the inhibition of the PCR reaction. Diluting the cDNA samples to overcome the inhibition was proven successful by conventional PCR (assessed by loading the PCR product in an agarose gel) but the cDNA concentration after dilution was not enough for RT-qPCR amplification. The tested primers and thermal programs had been successful in amplifying *nirK* in Cluster II of *F. imtechensis* obtained after cloning and for that reason 6 ng of DNA from microcosms samples was analysed for *nirK* relative expression. To quantify absolute numbers of copies PCR products of *nirS* and *nosZ* gene were obtained as described above for *R. denitrificans*. Additionally, *R. denitrificans* was also used to obtain PCR products of the *16S rRNA* gene using the primers described in Table S2 and thermocycler conditions described in a previous work.^19^ PCR products were cloned using the TOPO-TA cloning kit with pCR 2.1-TOPO and One Shot TOP10 Chemically Competent *E. coli* (Invitrogen – Thermo Fisher). Colonies developed for 10 h in solid medium (40 g L^-1^ LB Broth, ampicillin 50 μg ml^-1^ and X-gal 100 mmol L^-1^) at 37 °C, and then selected plasmid-inserted colonies were grown overnight in liquid medium. Plasmids were isolated from this culture using GenElute Plasmid MiniPrep kit (Sigma-Aldrich). Confirmation of the correct gene fragments was done by Sanger sequencing and genes were amplified in a StepOnePlus real-time PCR System, as described above. A standard curve was constructed by plotting the quantification cycle (Cq) versus the logarithm of the number of copies of the gene, assuming for standard DNA 660 g mol^-1^ and 6.022 x 10^23^ bp mol^-1^.^20^ A correction for ploidy was introduced for *16S rRNA* following a search in *rrn*DB database^21^ and considering an average of three *16S rRNA* operon copy number per genome. For *nirS* and *nosZ* a conservative approach (1 copy per organism) was considered, following a previous work listing organisms harbouring denitrification genes, with the respective copy numbers.^12^

### Supplementary Tables

**Table S1.** Predicted OTU-containing *nir* and *nosZ* on the sediment microbial community of Douro estuary.

| Clade | OTU | Kingdom | Phylum | Class | Order | Family | Genus | 2014<br>Jun | 2015<br>Jun | 2014<br>Dec | 2015<br>Dec |
| --- | --- | --- | --- | --- | --- | --- | --- | --- | --- | --- | --- |
| *nirK* I | C2_1511 | Bacteria | Proteobacteria | Rhizobiales | Bradyrhizobiaceae | Alphaproteobacteria | Bradyrhizobium | 0.00000 | 0.00000 | 0.00998 | 0.00000 |
| *nirK* I | C2_4712 | Bacteria | Proteobacteria | Rhizobiales | Bradyrhizobiaceae | Alphaproteobacteria | Bradyrhizobium | 0.00000 | 0.00000 | 0.00998 | 0.00249 |
| *nirK* II | C2_975 | Bacteria | Proteobacteria | Burkholderiales | Comamonadaceae | Betaproteobacteria | Acidovorax | 0.08072 | 0.11641 | 0.44897 | 0.23670 |
| *nirS* I | A1_10994 | Bacteria | Proteobacteria | Pseudomonadales | Pseudomonadaceae | Gammaproteobacteria | Pseudomonas | 0.01883 | 0.03004 | 0.00998 | 0.01246 |
| *nirS* I | A1_13628 | Bacteria | Proteobacteria | Pseudomonadales | Pseudomonadaceae | Gammaproteobacteria | Pseudomonas | 0.01883 | 0.04506 | 0.03991 | 0.05481 |
| *nirS* I | A1_1863 | Bacteria | Proteobacteria | Rhodocyclales | Rhodocyclaceae | Betaproteobacteria | Azoarcus | 0.05919 | 0.03380 | 0.04989 | 0.06229 |
| *nirS* I | A1_2344 | Bacteria | Proteobacteria | Pseudomonadales | Pseudomonadaceae | Gammaproteobacteria | Pseudomonas | 0.02153 | 0.03755 | 0.01995 | 0.02990 |
| *nirS* I | A1_7747 | Bacteria | Proteobacteria | Rhodocyclales | Rhodocyclaceae | Betaproteobacteria | Azoarcus | 0.06727 | 0.07886 | 0.14966 | 0.09966 |
| *nirS* I | A1_827 | Bacteria | Proteobacteria | Pseudomonadales | Pseudomonadaceae | Gammaproteobacteria | Pseudomonas | 0.82333 | 0.87123 | 1.27706 | 1.16604 |
| *nirS* I | C2_1511 | Bacteria | Proteobacteria | Rhizobiales | Bradyrhizobiaceae | Alphaproteobacteria | Bradyrhizobium | 0.00000 | 0.00000 | 0.00998 | 0.00000 |
| *nirS* I | C2_4712 | Bacteria | Proteobacteria | Rhizobiales | Bradyrhizobiaceae | Alphaproteobacteria | Bradyrhizobium | 0.00000 | 0.00000 | 0.00998 | 0.00249 |
| *nirS* I | C2_9 | Bacteria | Proteobacteria | Rhodocyclales | Rhodocyclaceae | Betaproteobacteria | Dechloromonas | 0.00538 | 0.00751 | 0.00998 | 0.01495 |
| *nirS* I | DA2_28640 | Bacteria | Proteobacteria | Pseudomonadales | Pseudomonadaceae | Gammaproteobacteria | Pseudomonas | 0.01345 | 0.02253 | 0.01995 | 0.05980 |
| *nosZ* I | A1_10994 | Bacteria | Proteobacteria | Pseudomonadales | Pseudomonadaceae | Gammaproteobacteria | Pseudomonas | 0.01883 | 0.03004 | 0.00998 | 0.01246 |
| *nosZ* I | A1_13628 | Bacteria | Proteobacteria | Pseudomonadales | Pseudomonadaceae | Gammaproteobacteria | Pseudomonas | 0.01883 | 0.04506 | 0.03991 | 0.05481 |
| *nosZ* I | A1_1635 | Bacteria | Proteobacteria | Alteromonadales | Alteromonadaceae | Gammaproteobacteria | Marinobacter | 0.15606 | 0.13519 | 0.16961 | 0.20181 |
| *nosZ* I | A1_1863 | Bacteria | Proteobacteria | Rhodocyclales | Rhodocyclaceae | Betaproteobacteria | Azoarcus | 0.05919 | 0.03380 | 0.04989 | 0.06229 |
| *nosZ* I | A1_2344 | Bacteria | Proteobacteria | Pseudomonadales | Pseudomonadaceae | Gammaproteobacteria | Pseudomonas | 0.02153 | 0.03755 | 0.01995 | 0.02990 |
| *nosZ* I | A1_23555 | Bacteria | Proteobacteria | Rhodobacterales | Rhodobacteraceae | Alphaproteobacteria | Roseovarius | 0.08879 | 0.11266 | 0.07982 | 0.10464 |
| *nosZ* I | A1_25707 | Bacteria | Proteobacteria | Rhodobacterales | Rhodobacteraceae | Alphaproteobacteria | Roseovarius | 0.03767 | 0.02629 | 0.01995 | 0.03737 |
| ***nosZ I*** | A1_29855 | Bacteria | Proteobacteria | Rhodobacterales | Rhodobacteraceae | Alphaproteobacteria | Roseovarius | 0.01883 | 0.07135 | 0.04989 | 0.04236 |
| ***nosZ I*** | A1_7747 | Bacteria | Proteobacteria | Rhodocyclales | Rhodocyclaceae | Betaproteobacteria | Azoarcus | 0.06727 | 0.07886 | 0.14966 | 0.09966 |
| ***nosZ I*** | A1_7862 | Bacteria | Proteobacteria | Alteromonadales | Alteromonadaceae | Gammaproteobacteria | Marinobacter | 0.01614 | 0.01127 | 0.00000 | 0.00747 |
| ***nosZ I*** | A1_788 | Bacteria | Proteobacteria | Rhodobacterales | Rhodobacteraceae | Alphaproteobacteria | Roseovarius | 0.08610 | 0.14270 | 0.19954 | 0.09468 |
| ***nosZ I*** | A1_827 | Bacteria | Proteobacteria | Pseudomonadales | Pseudomonadaceae | Gammaproteobacteria | Pseudomonas | 0.82333 | 0.87123 | 1.27706 | 1.16604 |
| ***nosZ I*** | C2_1511 | Bacteria | Proteobacteria | Rhizobiales | Bradyrhizobiaceae | Alphaproteobacteria | Bradyrhizobium | 0.00000 | 0.00000 | 0.00998 | 0.00000 |
| ***nosZ I*** | C2_4712 | Bacteria | Proteobacteria | Rhizobiales | Bradyrhizobiaceae | Alphaproteobacteria | Bradyrhizobium | 0.00000 | 0.00000 | 0.00998 | 0.00249 |
| ***nosZ I*** | C2_7831 | Bacteria | Proteobacteria | Rhodocyclales | Rhodocyclaceae | Betaproteobacteria | Thauera | 0.00269 | 0.00000 | 0.00000 | 0.00000 |
| ***nosZ I*** | C2_975 | Bacteria | Proteobacteria | Burkholderiales | Comamonadaceae | Betaproteobacteria | Acidovorax | 0.08072 | 0.11641 | 0.44897 | 0.23670 |
| ***nosZ I*** | DA1_5583 | Bacteria | Proteobacteria | Rhodobacterales | Rhodobacteraceae | Alphaproteobacteria | Roseovarius | 0.02960 | 0.04506 | 0.02993 | 0.04485 |
| ***nosZ I*** | DA2_28640 | Bacteria | Proteobacteria | Pseudomonadales | Pseudomonadaceae | Gammaproteobacteria | Pseudomonas | 0.01345 | 0.02253 | 0.01995 | 0.05980 |
| ***nosZ II*** | A1_116 | Bacteria | Bacteroidetes | Flavobacteriales | Flavobacteriaceae | Flavobacteriia | Maribacter | 0.15337 | 0.16523 | 0.14966 | 0.15447 |
| ***nosZ II*** | A1_4166 | Bacteria | Bacteroidetes | Flavobacteriales | Flavobacteriaceae | Flavobacteriia | Maribacter | 0.04305 | 0.02629 | 0.01995 | 0.05980 |

Data is from a previous survey at Douro estuary<sup>11</sup> and corresponds to four samplings at the same site in December and June of 2014 and 2015; taxonomic classification was obtained with SILVA SSU Ref dataset (v128; http://www.arb-silva.de).

**Table S2.**
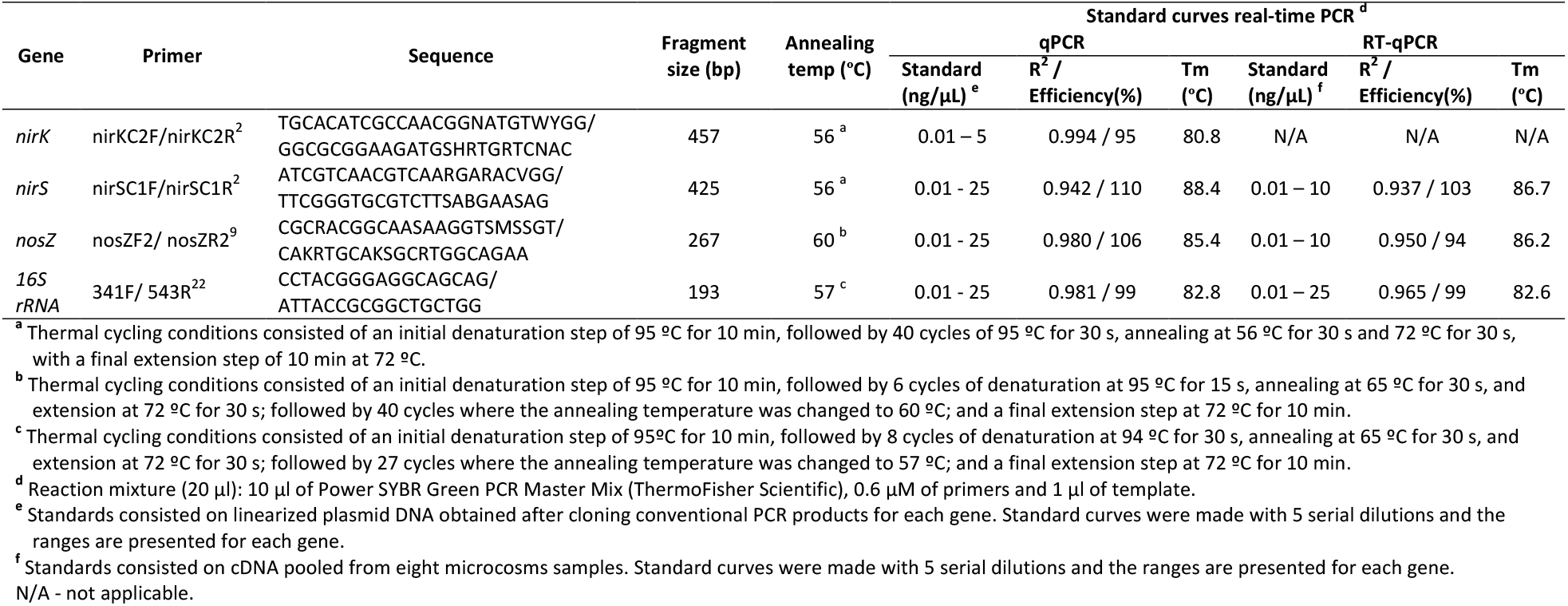
Real-time PCR conditions and respective sequencing primers

**Table S3.** *p*-values of the regressions over time for microcosms exposed to Cu NPs < 50 nm (EXP50) and exposed to Cu NPs < 150 nm (EXP150), and *p*-values for the *t*-test comparing the slopes of both EXP

|  | **pH** | **N-NO<sub>3</sub><sup>-</sup> water** |
| --- | --- | --- |
| **EXP50** | 1.68E-04 | 9.59E-07 |
| **EXP150** | 1.04E-03 | 1.63E-02 |
| **EXP50 vs EXP150** | 6.86E-01 | 7.40E-05 |

### Supplementary Figures

**Fig. 1S.**
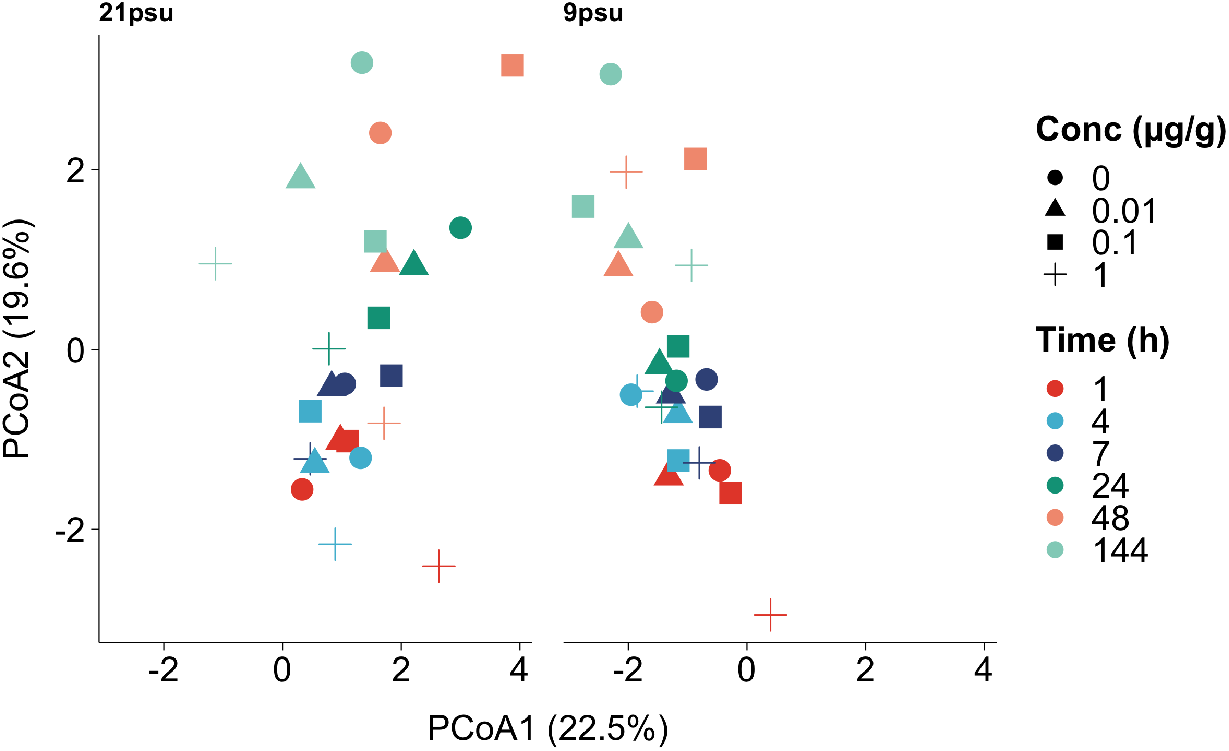
Microcosms response throughout the exposure period under different salinities (9 psu and 21 psu). Principal component analysis showing the Cu concentration throughout time to behave similarly in microcosms exposed to Cu NPs < 50 nm under different salinity regimes.

**Fig. 2S.**
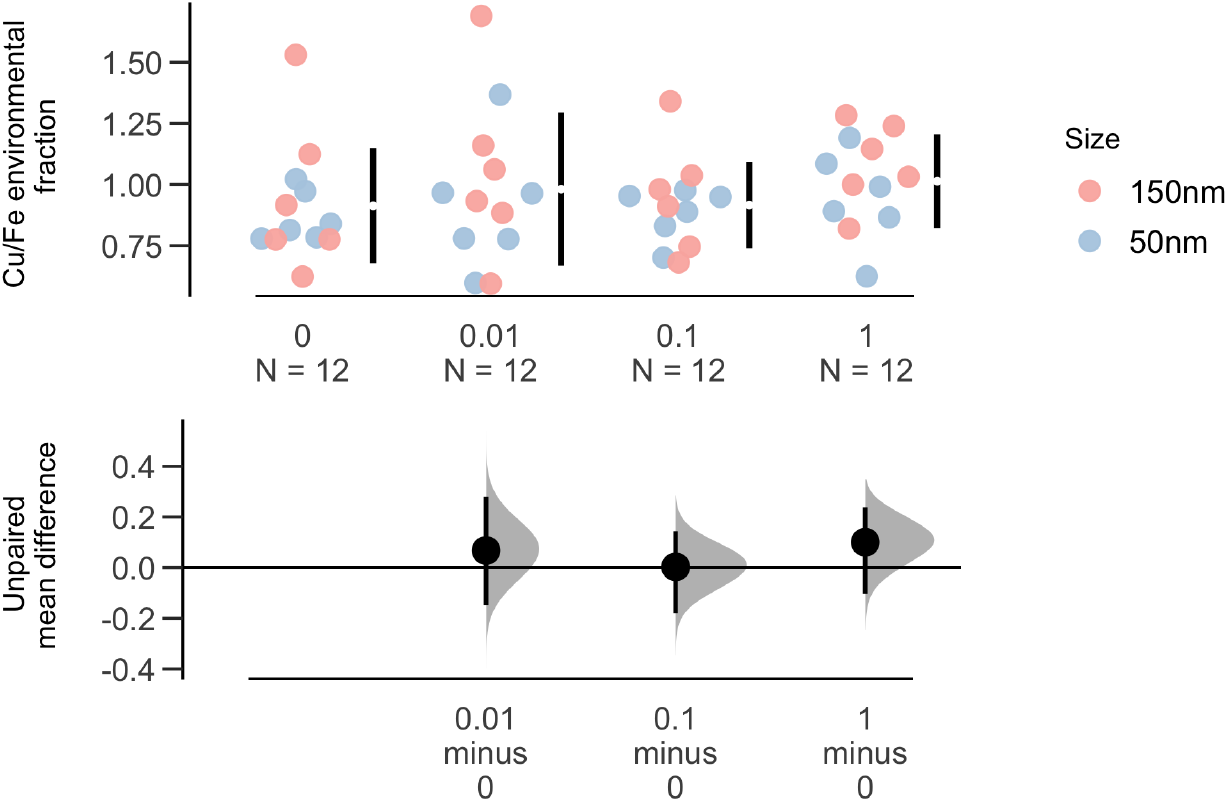
Estimation plot of Cu environmental fraction normalised to Fe. Results are reported for microcosms exposed to Cu NPs 0, 0.01, 0.1 and 1 μg g^-1^. Gapped lines to the right of each concentration group are mean value ± standard deviation and the effect size, with 95 % confidence intervals, is plotted below.

**Fig. 3S.**
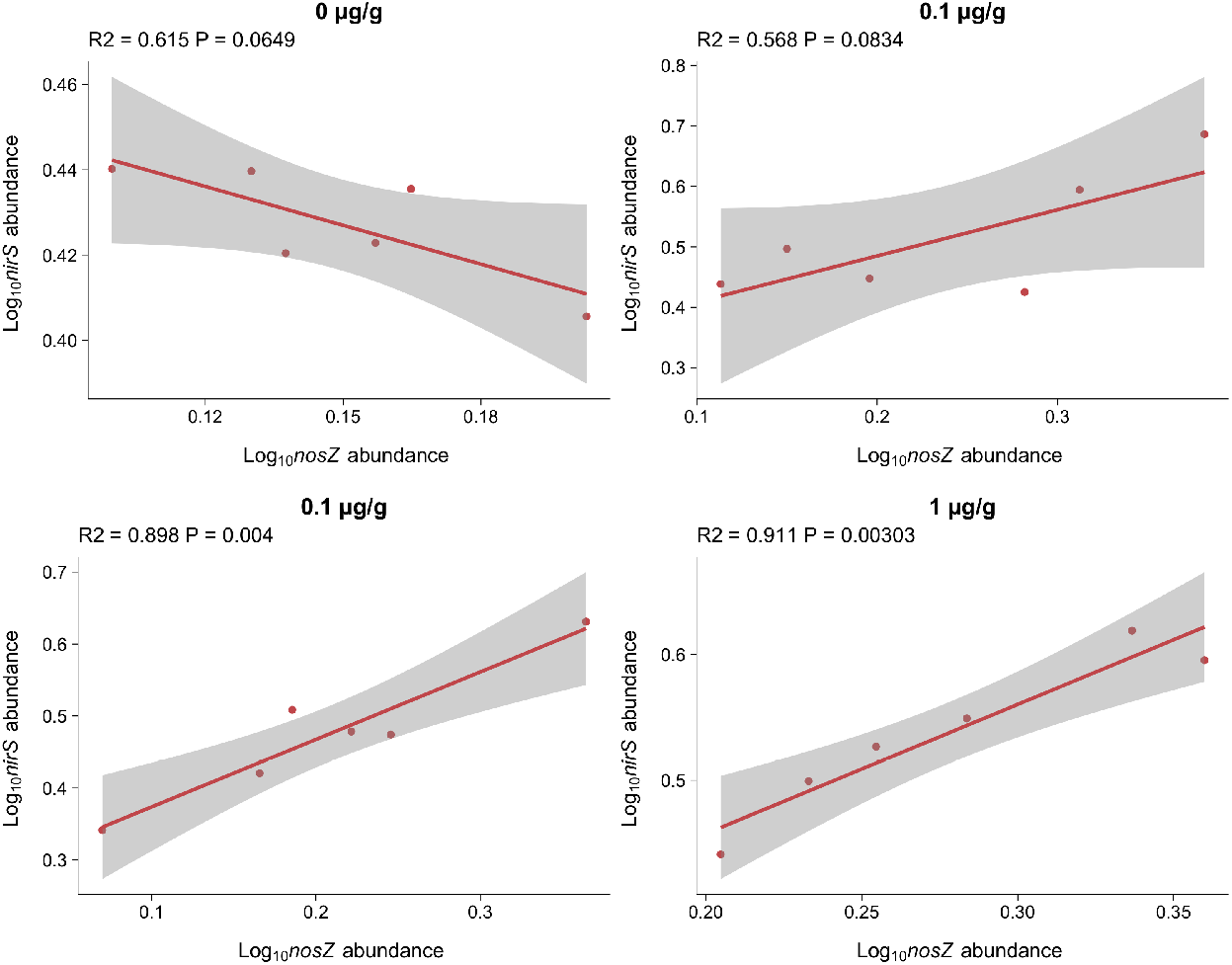
Correlation between *nirS* and *nosZ* abundances in microcosms exposed to Cu NPs < 50 nm. **A)** microcosms not exposed to Cu NPs; **B)** microcosms exposed to 0.01 μg g^-1^; **C)** microcosms exposed to 0.1 μg g’^1^; and **D)** microcosms exposed to 1 μg g^-1^. Gene copy number per gram dry sediment were log10 transformed and *nirS* and *nosZ* values were normalized by the *16S rRNA* gene. Points are the average of two replicates, with a RSD between 9 and 24 % and the linear regression is shown with 95 % confidence interval.

## Notes

### Competing Interest Statement

The authors have declared no competing interest.

